# Human DC3 Antigen Presenting Dendritic Cells from Induced Pluripotent Stem Cells

**DOI:** 10.1101/2021.06.22.449451

**Authors:** Taiki Satoh, Marcelo A. S. Toledo, Janik Boehnke, Kathrin Olschok, Niclas Flosdorf, Katrin Götz, Caroline Küstermann, Stephanie Sontag, Kristin Seré, Steffen Koschmieder, Tim H. Brümmendorf, Nicolas Chatain, Yoh-ichi Tagawa, Martin Zenke

**Author notes:** **Correspondence:** Martin Zenke, PhD, Institute for Biomedical Engineering, Department of Cell Biology, RWTH Aachen University Hospital, Pauwelsstrasse 30, 52074 Aachen, Germany.

## Abstract

Dendritic cells (DC) are professional antigen-presenting cells that develop from hematopoietic stem cells. Different DC subsets exist based on ontogeny, location and function, including the recently identified proinflammatory DC3 subset. DC3 have the prominent activity to polarize CD8^+^ T cells into CD8^+^ CD103^+^ tissue resident T cells. Here we describe human DC3 differentiated from induced pluripotent stem cells (iPS cells). iPS cell-derived DC3 have the gene expression and surface marker make-up of blood DC3 and polarize CD8+ T cells into CD8+ CD103+ tissue-resident memory T cells in vitro. To test the impact of malignant JAK2 V617F mutation on DC3, we differentiated patient-specific iPS cells with JAK2 V617F^het^ and JAK2 V617F^hom^ mutations into JAK2 V617F^het^ and JAK2 V617F^hom^ DC3. The JAK2 V617F mutation enhanced DC3 production and caused a bias towards erythrocytes and megakaryocytes. The patient-specific iPS cell-derived DC3 are expected to allow studying DC3 in human diseases and developing novel therapeutics.

**Graphical Abstract:** 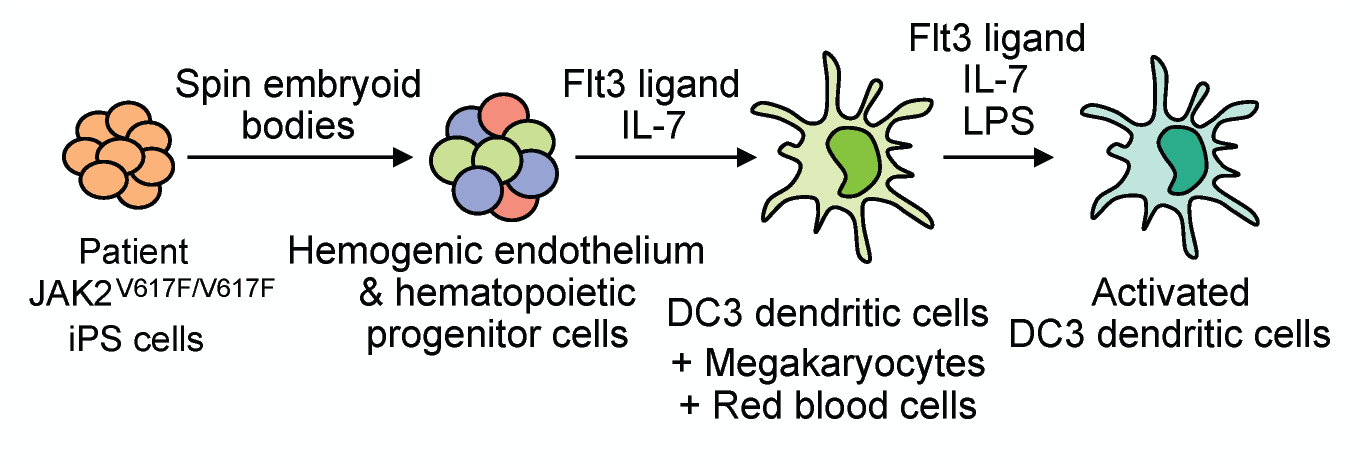

## Introduction

Dendritic cells (DC) develop from hematopoietic stem cells and populate a large array of lymphoid and non-lymphoid tissues in our body. DC play a pivotal role in antigen presentation to induce immunity and immunological tolerance (Guilliams et al., 2014; Mildner and Jung, 2014). DC are classified into two main subsets, classical DC (cDC) and plasmacytoid DC (pDC), based on surface marker expression, function and ontogeny, and cDC are further subdivided into cDC1 and cDC2 (Guilliams et al., 2014; Mildner and Jung, 2014; Collin and Bigley, 2018; Amon et al., 2020). However, DC heterogeneity and in particular cDC2 heterogeneity have been controversial to date. Single cell techniques revealed two subpopulation in cDC2 referred to as DC2 and DC3 (Villani et al., 2017; Brown et al., 2019; Dutertre et al., 2019). DC3 share features with both DC2 and monocytes but are developmentally and functionally different from both DC2 and monocytes (Bourdely et al., 2020; Cytlak et al., 2020). All cDC subsets activate T cells but DC3 have the prominent activity to polarize CD8^+^ T cells into CD8^+^ CD103^+^ tissue-resident memory (T_RM_) T cells (Bourdely et al., 2020). DC are a scarce cell type in blood and tissues (Granot et al., 2017), which hampers their analysis in mice and man, and their therapeutic application in the human system.

Pluripotent stem cells, including embryonic stem cells (ES cells) and induced pluripotent stem cells (iPS cells), are a particular appealing cell source for studies in developmental biology and disease modeling, and in the human system for regenerative medicine (Rowe and Daley, 2019). Patient and disease-specific iPS cells capture disease-specific and/or associated mutations and this is key to model human diseases in a dish and for compound screening. Human ES cells and iPS cells can be induced to differentiate into hematopoietic cells, often through 3D dimentional embryoid body (EB) differentiation (Sturgeon et al., 2014; Ivanovs et al., 2017; Garcia-Alegria et al., 2018), and many blood cell types were generated from human iPS cells, including DC (Ackermann et al., 2015). Frequently, the cytokine cocktails to induce DC development included GM-CSF (Senju et al., 2011; Silk et al., 2012; Cai et al., 2017; Sontag et al., 2017; Sachamitr et al., 2018; Horton et al., 2020), yet DC development is known to dependent on Flt3 ligand (Flt3L) (Felker et al., 2010; Mildner and Jung, 2014; Collin and Bigley, 2018; Amon et al., 2020).

The JAK2 V617F mutation is associated with a group of hematopoietic malignancies termed myeloproliferative neoplasms (MPN), including polycythemia vera (PV), essential thrombocythemia (ET) and primary myelofibrosis (PMF) (Vainchenker and Kralovics, 2017). JAK2 V617F causes cytokine-independent constitutive activation of JAK2 kinase and its signaling pathways, thereby leading to aberrant production of red cells, platelets and myeloid cells in MPN. Whether this involves also an aberrant regulation of immune cells, such as DC, and how this might cause and/or contribute to disease is largely unknown.

To investigate the impact of proinflammatory JAK2 V617F signaling on DC we differentiated disease-specific JAK2 V617F iPS cells into DC. Frequently, DC are generated in vitro with GM-CSF, which triggers the JAK2/STAT signaling pathway (Hacker et al., 2003; Hieronymus et al., 2005; Seré et al., 2012; Collin and Bigley, 2018; Zhan et al., 2019) and thus a high dose of GM-CSF stimulation might hide the activity of JAK2 V617F mutation in DC. Here we differentiated human JAK2 V617F iPS cells into DC using Flt3L and IL-7. JAK2 V617F mutation did not affect DC function but enhanced production of DC, erythrocytes and megakaryocytes. Interestingly, the CD1c^+^ DC obtained exhibited properties of DC3 and polarized CD8^+^ T cells towards CD8^+^ CD103^+^ T_RM_ cells.

## Materials and Methods

### Generation and culture of patient-specific iPS cells with JAK2 V617F mutation

Patient-specific iPS cells were obtained by reprogramming peripheral blood mononuclear cells (PBMC) of healthy donor and of two PV patients (referred to as PV1 and PV2) with JAK2 V617F mutation after informed consent (local board reference number EK099/14 and EK206/09) with OCT4, SOX2, c-MYC and KLF4 in CytoTune Sendai virus vectors (Thermo Fisher Scientific) (Sontag et al., 2017). iPS cells with monoallelic and biallelic JAK2 V617F mutation, referred to as JAK2 V617F^het^ and JAK2 V617F^hom^ iPS cells, respectively, and without mutation were isolated and further studied. Patient PV2 exhibited a high JAK2 V617F allele burden (96%) and only JAK2 V617F^hom^ iPS cell clones were obtained, and thus the isogenic unmutated control was generated by CRISPR/Cas9 editing using the Alt-R CRISPR/Cas9 system (IDT, Coralville, USA).

Briefly, the CRISPR/Cas9 complex (Alt-R HiFi Cas9 nuclease plus gRNA), single-stranded donor template and electroporation enhancer were delivered to cells using the Neon Transfection System and the 100 μl kit (Thermo Fisher Scientific). Before electroporation, iPS cells were treated for 1 h with HDR enhancer (5 μM; IDT, Coralville, USA) and 10 μM Y-27632 (Abcam, Cambridge, UK). Electroporated cells were seeded on Laminin 521 (Biolamina, Sundbyberg, Sweden) coated plates in StemMACS iPS-Brew XF (Miltenyi Biotec) supplemented with 1x CloneR (Stemcell Technologies, Vancouver, Canada). Genotyping of CRISPR-repaired iPS cell lines was performed by allele-specific PCR targeting the JAK2 V617F mutation. Sequences of crRNA, donor template and allele-specific PCR primers are listed in Supplementary Table 1. Potential off-target genes CREBL2 and COA6 were unaffected (Supplementary Figure 1).

In the Human Pluripotent Stem Cell Registry (www.hpscreg.eu) PV1 JAK2 and JAK2 V617F^het^ iPS cells are referred as UKAi002-A and UKAi002-B, and PV2 JAK2 and JAK2 V617F^hom^ iPS cells are referred to as UKAi003-A2 and UKAi003-A, respectively.

Healthy donor iPS cells were obtained by reprogramming PBMC as above and were used as healthy control. Routinely, iPS cells were maintained in StemMACS iPS-Brew XF (Miltenyi Biotec) on 6-well plates coated with Matrigel (Corning).

### iPS cell differentiation into hematopoietic cells and DC

Differentiation of human iPS cells into hemogenic endothelium (HE) and hematopoietic progenitor cells (HPC) was by spin embryoid bodies (spin EB) protocol (Liu et al., 2015) with some modifications. Briefly, iPS cells were harvested with Accutase and single cell suspension were plated on round-bottom 96 well suspension plates (Greiner) with 4000 cells/well in serum free medium (SFM) containing 10 ng/ml BMP4 (Miltenyi Biotec), 10 ng/ml bFGF (Peprotech), 10 μM Y-27632 (Abcam), 50 μg/ml L-ascorbic acid (L-AA, Stem Cell Technologies) and 6 μg/ml holo-transferrin (Sigma). SFM was a 1:1 mixture of IMDM and F12 medium (both Thermo Fisher Scientific) containing 0.5% BSA, 2 mM GlutaMAX, 1% chemically defined lipid concentrate (both Thermo Fisher Scientific) and 400 μM 1-thioglycerol (MTG, Sigma). On day 2 fresh culture medium containing BMP4, bFGF, L-AA, holo-transferrin, and 10 ng/ml VEGF (Peprotech) was added.

EB formation was observed after 24-48 hours and from day 4 onwards half medium change was performed by adding fresh culture medium containing bFGF, VEGF, L-AA, holo-transferrin, and SCF (0.5% supernatant of SCF producing CHO KLS cells) every second day. On day 10-11 EB were harvested by gentle pipetting and 30 EB/well were plated on gelatin-coated 6-well plates in RPMI (Thermo Fisher Scientific), 10% FCS (PAN Biotech) with 25 ng/ml Flt3L (Peprotech), 10 ng/ml IL-7 (Miltenyi Biotec), 2 mM L-Glutamine, and 100 μM 2-Mercaptoethanol (both Thermo Fisher Scientific) referred to as DC culture medium. Four days later fresh DC culture medium was added and on day 8-9 of differentiation suspension and loosely adherent cells were harvested with gentle pipetting and HLA-DR^+^ cells were isolated by immunomagnetic bead selection (MACS) with HLA-DR microbeads (Miltenyi Biotec) following the manufacture’s instruction. HLA-DR^+^ cells were resuspended in DC culture medium without or with 1 μg/ml LPS and cultured for one day to induce the DC activation. These HLA-DR^+^ cells were used for further experiments.

### Flow cytometry analysis and Diff-Quik staining

Hemogenic endothelium (HE, CD34+ CD31+ CD144+ CD43- CD45- CD73-) hematopoietic progenitor cells (HPC, CD43^+^ CD34^low/^-), cells obtained upon DC differentiation and T cells were analyzed by flow cytometry (Supplementary Figure 2A-E). Cells were stained with specific antibodies (Supplementary Table 2) and analyzed on FACS Canto II (BD) as described (Sontag et al., 2017) and data were analyzed with FlowJo software (Tree Star).

For Diff-Quik staining HLA-DR^+^ cells were centrifuged onto glass slides in Cytospin 4 cytocentrifuge (Thermo Fisher Scientific). Cells were fixed with methanol, stained with Diff-Quik (Medion Dianostics) and mounted with Entellan (Merck).

### Gene expression analysis by RT-qPCR

Total RNA was isolated with NucleoSpin RNA Kit (Macherey Nagel) following the manufacturer’s instruction, quantified with NanoDrop (Thermo Fisher Scientific) and subjected to reverse transcription-quantitative polymerase chain reaction (RT-qPCR) analysis. Briefly, total RNA was reverse transcribed with High capacity cDNA Reverse Transcriptase Kit (Thermo Fisher Scientific). Synthesized cDNA was used for qPCR analysis with FAST SYBR Green master mix (Applied BioSystem) on a StepOne Plus device (Applied BioSystem). Expression values were normalized to *GAPDH* and ΔCt value over 12 was regarded as not-expressed and arbitrarily set to 12. z-scores of -ΔCt mean were calculated for each gene and subjected to hierarchical clustering and representation in heat map format with Morpheus software (https://software.broadinstitute.org/morpheus/). Primers are listed in Supplementary Table 3.

### Chemotaxis assay

Chemotaxis assay was performed as described (Kurz et al., 2002; Sontag et al., 2017). In brief, transwell inserts (5 μm pore size, Corning) were incubated with culture medium for 1 hour (37°C, 5% CO2) to block unspecific binding. In the lower chamber, medium with or without 100 ng/ml CCL19 (Peprotech) was added and then 3-4 × 10^4^ HLA-DR^+^ cells were added to the upper chamber. After 2 h incubation (37°C, 5% CO_2_), 10^4^ Dynabeads (15 μm diameter, Dynal polymers) were added to the lower chamber, to allow normalization for variations in the experimental procedure, and cells and beads of the lower chamber were analyzed by flow cytometry as described before (Kurz et al., 2002). The number of migrated cells was determined relative to 10^4^ beads from the ratio of cells/beads analyzed by flow cytometry (Supplementary Figure 2D) and the percentage of migrated cells was calculated relative to the total input number of cells.

### Mixed lymphocyte reaction (MLR)

MLR was performed as described (Dutertre et al., 2019) with some modifications. PBMC were obtained from healthy donors after informed consent. CD3^+^ T cells were isolated by MACS with CD3 microbeads (Miltenyi Biotec) and labeled with 5 μM CFSE (Stem Cell Technologies) for 20 min at 37°C. CFSE-labeled CD3^+^ T cells (10^5^ cells) were mixed with unstimulated or LPS-stimulated HLA-DR^+^ cells (10^4^ cells) and cultured in 10% KO-serum replacement IMDM (Thermo Fisher Scientific) for 5 days. CFSE-labeled CD3^+^ T cells were cultured with 5-10 μg/ml concanavalin A (Con A) or without Con A, to provide positive and negative controls, respectively. Before harvesting the cells, 10^4^ Dynabeads were added and CFSE division and T cell numbers were analyzed by flow cytometry and normalized to beads.

### Statistical Analysis

Statistical analyses were performed in Prism 8 (GraphPad) using one-way or two-way ANOVA with tukey’s multiple comparisons test or uncorrected fisher’s LSD test. Results were considered to be significant at p < 0.05 (*), p < 0.005 (**), p < 0.001 (***), p < 0.0001 (****) with specific comparisons as indicated in respective figure legends.

## Results

### Human iPS cells differentiate into CD1c^+^ DC with DC3 characteristics

Human iPS cells were differentiated into HE and HPC in a spin EB protocol (Figure 1A). To establish iPS cell differentiation we employed iPS cells of healthy donor and on day 10 of EB differentiation CD34^+^ CD43- HE and CD43^+^ CD34^low/-^ HPC were observed (Figure 1B). EB were harvested and further differentiated into DC with Flt3L and IL-7, thereby following a protocol modified from DC differentiation of cord blood cells (Balan et al., 2018; Laustsen et al., 2018). Monitoring the kinetics of major histocompatibility class II (MHC II) expression by flow cytometry revealed a peak of CD1c^+^ HLA-DR^+^ cells on day 8 of differentiation (data not shown). About 40% HLA-DR^+^ cells were routinely obtained on day 8-9 (Figure 1C) and HLA-DR^+^ cells were isolated and used for further analyses (Figure 1A).

**Figure 1.**
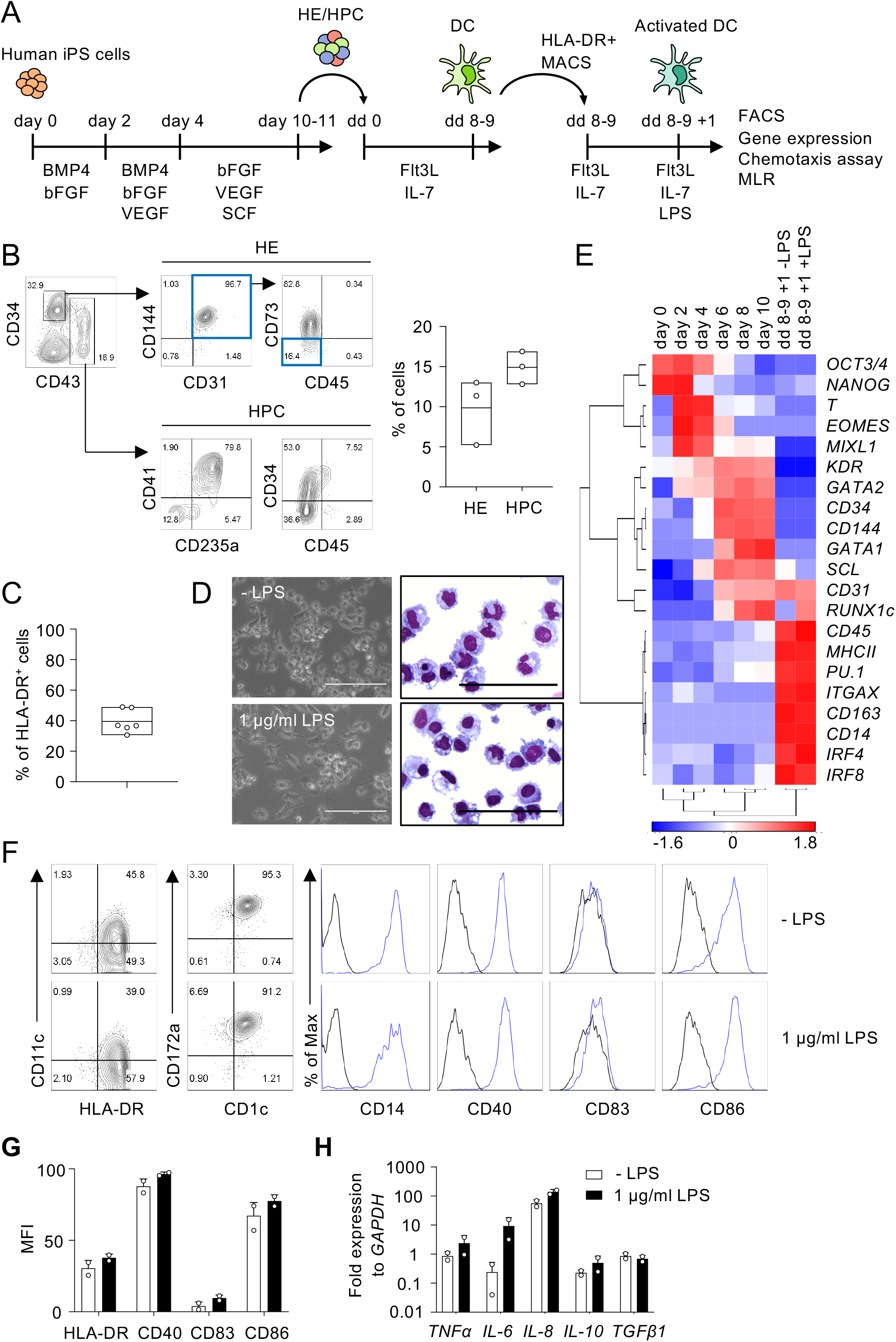
Human iPS cells differentiate into DC in EB and Flt3L + IL-7 cultures. **(A)** Schematic representation of iPS cell differentiation into DC. Human iPS cells were differentiated into HE and HPC for 10-11 days in spin EB culture with consecutive supplementation of cytokines as indicated. HE and HPC were then cultured for 8-9 days with Flt3 ligand (Flt3L) and IL-7 to induce DC differentiation (dd 0 to dd 8-9). HLA-DR^+^ cells were isolated by immunomagnetic bead selection (MACS) and stimulated with 1 μg/ml LPS for one day (dd 8-9 +1). These cells were used for further analysis as indicated. **(B)** Representative flow cytometry analysis and quantification of HE (CD34^+^ CD31^+^ CD144^+^ CD43- CD45- CD73- cells; blue boxes) and HPC (CD43^+^ CD34^low/-^ cells) in EB on day 10 of spin EB differentiation. n = 3, line represents mean. **(C)** Frequency of HLA-DR^+^ cells on day 8-9 of DC differentiation. n = 6, line represents mean. **(D)** Representative photomicrographs of unstimulated and LPS-stimulated HLA-DR^+^ cells in culture (left) and in cytospin preparations stained with Diff-Quik (right). n = 6, scale bars: 100 μm. **(E)** Gene expression profiling of iPS cells (day 0) and EB on day 2-10 of spin EB differentiation, and unstimulated and LPS-stimulated HLA-DR^+^ cells on day 8-9 +1 of DC differentiation by RT-qPCR analysis. Expression values were normalized to GAPDH and ΔCt value over 12 was regarded as not-expressed and arbitrarily set to 12. z-scores of -ΔCt mean calculated for each gene are shown in heat map format (red, high expression; blue, low expression). n = 2. **(F and G)** Representative flow cytometry analysis and quantification of median fluorescent intensity (MFI) of DC markers on unstimulated and LPS-stimulated CD1c^+^ HLA-DR^+^ cells. Blue lines and black lines in histograms represent stained and unstained cells, respectively. MFI values were normalized to unstained cells. Panel (G): n = 2, mean ± SD. **(H)** Cytokine expression in LPS-stimulated CD1c^+^ HLA-DR^+^ cells by RT-qPCR analysis. Unstimulated cells (-LPS) are shown as control. Values were normalized to GAPDH and 2-ΔCt values in log scale were shown. n = 2, mean ± SD. All data shown in panels (B) – (H) are from healthy control iPS cells. Data in panels (D) and (F) are representative for healthy control and patient PV1 and PV2 iPS cells. n = 2-6.

HLA-DR^+^ cells showed DC morphology and expression of DC specific genes, such as *ITGAX* (*CD11c*), *MHCII*, *IRF4* and *IRF8*, and the expression of the DC3 genes *CD14* and *CD163* (Figure 1D and E). We then proceeded to investigate cDC subset specific markers, including CD141 and CLEC9A for cDC1, and CD1c and CD172a for cDC2. We also investigated CD163 and CLEC10A to distinguish DC and monocytes (Heger et al., 2018; Cytlak et al., 2020). The number of HLA-DR^+^ cells increased with time during DC differentiation and HLA-DR^+^ cells expressed CD1c and CD14 (Figure 1F, Supplementary Figure 3A). DC showed also expression of CD163, CD172a and CLEC10A, but not CD141 and CLEC9A, and (Figure 1F, Supplementary Figure 3B and Supplementary Figure 4A), implying that they are distinct from monocytes and resemble more DC3 than cDC1 or DC2.

DC expressed the co-stimulatory molecules CD40 and CD86, which relates to high HLA-DR expression in these cells, since they were enriched for HLA-DR expression (Figure 1A, F and G). There was no further up-regulation of HLA-DR, CD40, CD83, and CD86 expression upon stimulation with lipopolysaccharide (LPS; Figure 1F and G). DC expressed inflammatory cytokines, including *TNFα*, *IL-6* and *IL-8*, and there was essentially no further up-regulation upon LPS stimulation with the exception of some up-regulation of IL-6 (Figure 1H). Taken together, human iPS cells were efficiently induced to differentiate into DC with Flt3L and IL-7 and the CD1c^+^ HLA-DR^+^ DC obtained exhibited features of DC3.

### JAK2 V617F iPS cells differentiate into DC and JAK2 V617F enhances DC production

Next we applied this DC differentiation protocol to disease-specific iPS cells. JAK2 V617F iPS cells from two PV patients (hereafter referred to as PV1 and PV2) with monoallelic and biallelic JAK2 V617F mutation, referred to as JAK2 V617F^het^ and JAK2 V617F^hom^ iPS cells, respectively, and without mutation were studied. Due to the high JAK2 V617F allele burden in patient PV2, only clones with JAK2 V617F^hom^ mutation were obtained and thus the isogenic unmutated control was generated by CRISPR/Cas9 editing. JAK2 V617F^het^ and JAK2 V617F^hom^ iPS cells and unmutated controls were subjected to spin EB differentiation and CD34^+^ CD43- HE and CD43^+^ CD34^low/^- HPC were obtained after 10-11 days (Figure 2A). Cells were then differentiated into CD1c^+^ HLA-DR^+^ cells for 8-9 days. JAK2 V617F^het^ and JAK2 V617F^hom^ iPS cells produced more CD1c^+^ HLA-DR^+^ cells than iPS cells without mutation (Figure 2B). The frequency of CD1c^+^ HLA-DR^+^ cells for the different V617F genotypes of PV1 and PV2 varied (Supplementary Figure 4B), probably due to patient-specific differences and the increased erythropoiesis, megakaryopoiesis and granulopoiesis observed for these cells (see below Figure 3A and B).

**Figure 2.**
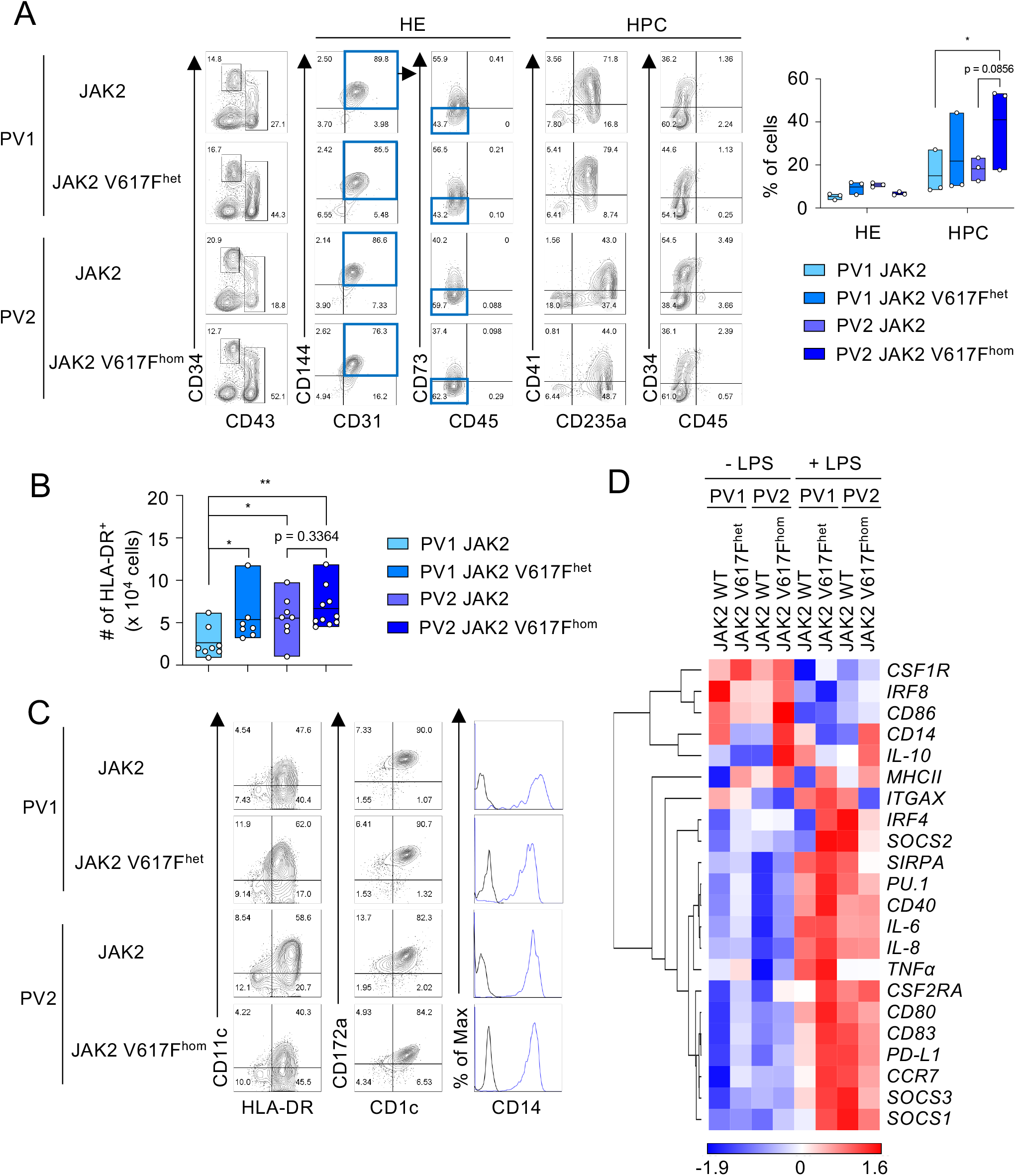
JAK2 V617F^het^ and JAK2 V617F^hom^ iPS cells differentiate into DC. **(A)** Representative flow cytometry analysis and quantification of HE and HPC (CD34^+^ CD31^+^ CD144^+^ CD43^-^ CD45^-^ CD73^-^ cells and CD43^+^ CD34^low/-^ cells, respectively) in EB on day 10 of spin EB differentiation. JAK2 V617F^het^ and JAK2 V617F^hom^ cells of PV patient 1 (PV1) and PV patient 2 (PV2), respectively, and cells of PV1 and PV2 without JAK2 V617F mutation (JAK2) are shown. n = 3, line represents mean; *p < 0.05, two-way ANOVA with tukey’s multiple comparisons test. **(B)** The number of HLA-DR^+^ cells on day 8-9 of DC differentiation. JAK2 genotypes are as in (A). n = 7-10, line represents mean; *p < 0.05, **p < 0.005, one-way ANOVA with uncorrected fisher’s LSD test. **(C)** Expression of the DC markers CD1c, CD11c and CD172a, and of CD14 on HLA-DR+ cells was assessed by flow cytometry. Blue lines and black lines in histograms show stained and unstained cells, respectively. **(D)** Gene expression profiling of unstimulated and LPS-stimulated CD1c^+^ HLA-DR^+^ cells on day 8-9 +1 of DC differentiation by RT-qPCR analysis. Expression values were normalized to GAPDH and z-scores of -ΔCt mean calculated for each gene are shown in heat map format as in Figure 1E. n = 3.

**Figure 3.**
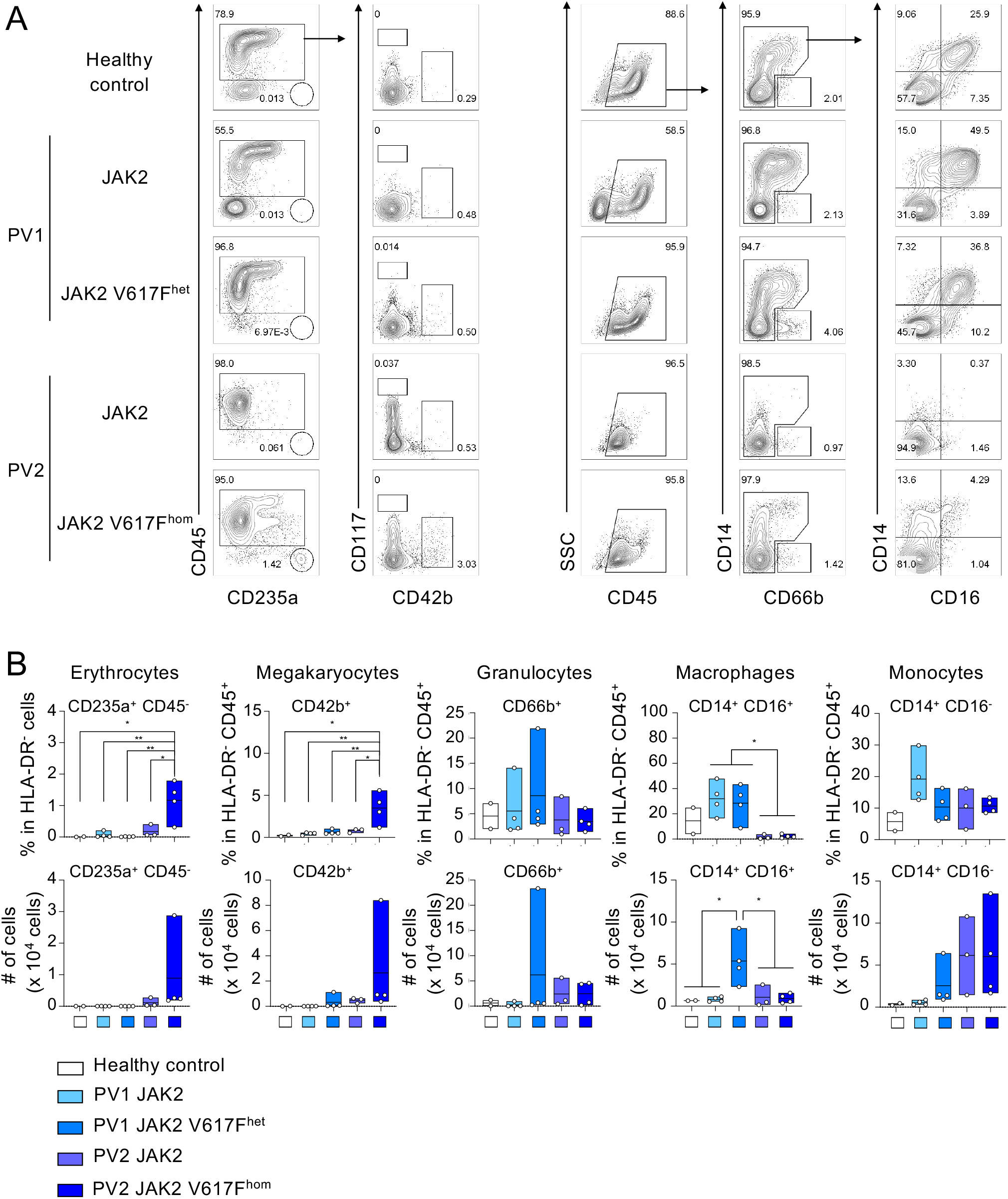
Erythropoiesis and megakaryopoiesis upon differentiation of JAK2 V617F^hom^ iPS cells. **(A)** JAK2 V617F^het^ and JAK2 V617F^hom^ iPS cells of patients PV1 and PV2, respectively, were differentiated as in Figure 1A and HLA-DR-cells on day 8-9 of differentiation were analyzed for erythrocytes, megakaryocytes and myeloid cells (granulocytes, monocytes and macrophages) by flow cytometry. Cells without mutation (JAK2) and cells of healthy donor (healthy control) were used as controls. Representative flow cytometry plots are shown. **(B)** Quantification of flow cytometry data of (A). n = 2-4, mean ± SD; *p < 0.05, **p < 0.005, oneway ANOVA with tukey’s multiple comparisons test.

Differentiated CD1c^+^ HLA-DR^+^ cells of all JAK2 V617F genotypes had the same DC phenotype based on cell morphology, surface maker and gene expression profiling and exhibited DC3 characteristics (Figure 2C and D, Supplementary Figure 4A). DC responded to LPS with augmented expression of proinflammatory and co-stimulatory molecules (Figure 2D, Supplementary Figure 4C and D).

To determine the impact of JAK2 V617F mutation on differentiation into other cell types than DC, HLA-DR^-^ cells of the DC differentiation protocol were analyzed. JAK2 V617F^hom^ caused a production of CD235a/glycophorin^+^ and CD42b^+^ cells (Figure 3A and B), indicative for erythropoiesis and megakaryopoiesis, respectively, which is in line with PV phenotype. A prominent CD14^+^ CD16 ^+^ macrophage population was observed for healthy control and patient PV1 but was essentially absent in patient PV2, again reflecting patient-specific differences.

In summary, the JAK2 V617F mutation enhanced production of CD1c^+^ HLA-DR^+^ DC without grossly affecting the DC phenotype and with preserving the DC3 characteristics.

### DC show chemotaxis towards CCL19 and polarize CD8^+^ T cells towards CD8^+^ CD103^+^ T_RM_ cells

Migration in response to chemokine gradient and T cell activation represent important DC properties. To this end, CD1c^+^ HLA-DR^+^ DC were activated with LPS or left untreated and subjected to chemotaxis toward CCL19. Both JAK2 V617F^hom^ DC and DC without mutation effectively migrated towards CCL19 chemokine (Figure 4A and Supplementary Figure 5A). JAK2 V617F^hom^ DC and unmutated controls also effectively induced CD4 and CD8 T cell proliferation (Figure 4B). DC3 polarize CD8^+^ T cells into CD8^+^ CD103^+^ T_RM_ cells (Bourdely et al., 2020), and thus we explored CD103 expression in proliferated T cells. JAK2 V617F^hom^ DC and unmutated DC induced CD103 expression in CD8 T cells, indicating polarization towards CD8^+^ CD103^+^ T_RM_ cells (Figure 4B). There was no CD103 expression in unstimulated T cells or T cells stimulated with concanavalin A (ConA). This T cell polarization is known to depend on TGFβ (Bourdely et al., 2020) and we note that the iPS cell-derived DC studied here show abundant expression of *TGFβ* (Supplementary Figure 5B).

**Figure 4.**
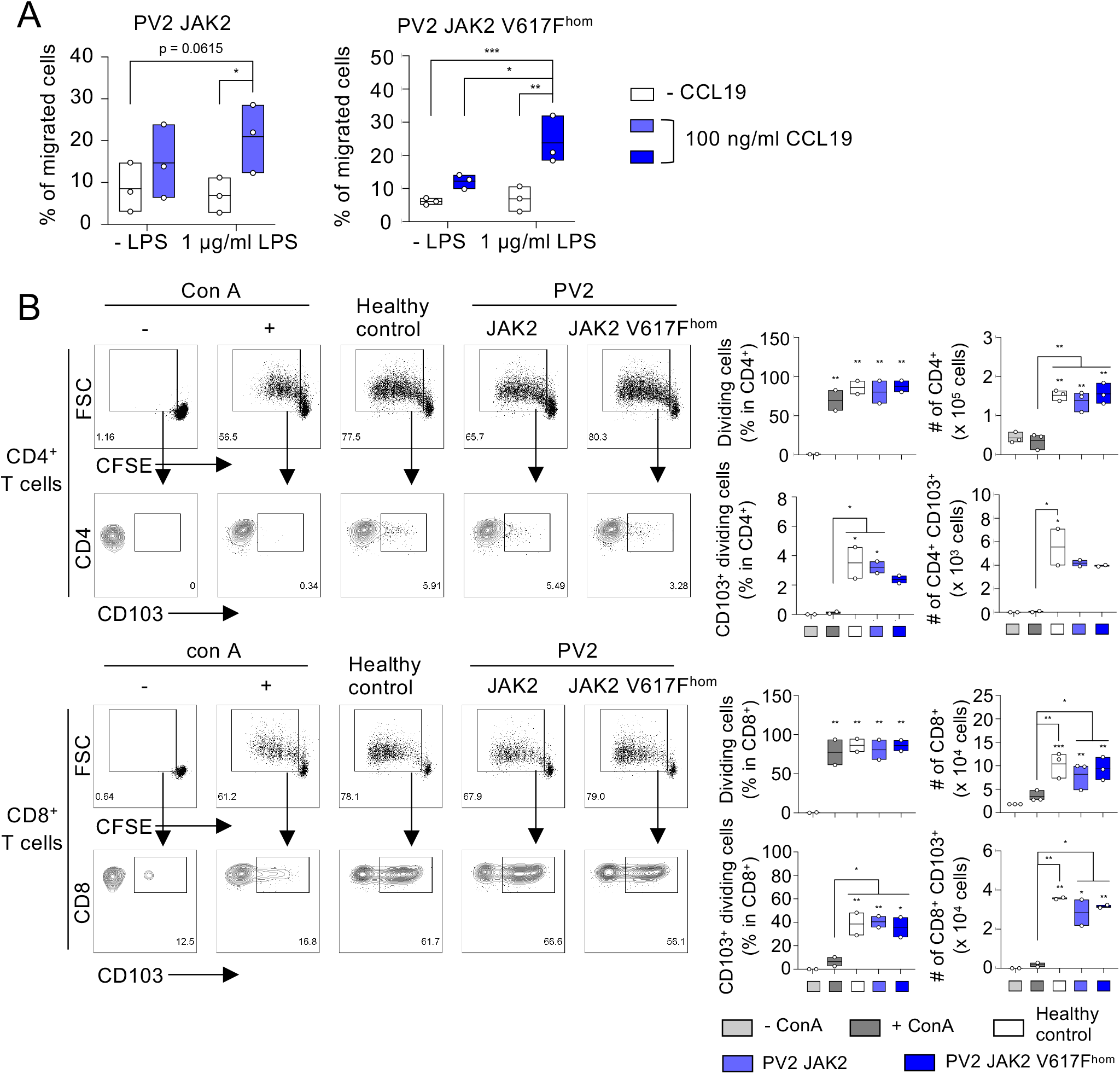
JAK2 V617F^hom^ DC migrate towards CCL19 chemokine and induce polarization of CD8+ T cells towards CD8+ CD103+ tissue resident T cells. **(A)** DC obtained from JAK2 V617F^hom^ iPS cells (patient PV2) and the respective iPS cells without mutation (JAK2) migrate towards CCL19 chemokine. CD1c^+^ HLA-DR^+^ cells of day 8-9 of DC differentiation were treated with LPS (1 μg/ml, 24 hours) or left untreated and subjected to CCL19 chemotaxis assay. Migrated cells in response to CCL19 were measured by flow cytometry and the percentage of migrated cells is shown. n = 3, line represents mean; *p < 0.05, **p < 0.005, two-way ANOVA with uncorrected fisher’s LSD test. **(B)** DC obtained from patient PV2 iPS cells as in (A) and from healthy donor iPS cells (healthy control) induce CD4^+^ and CD8^+^ T cell proliferation in MLR assays. CD1c^+^ HLA-DR^+^ cells on day 8-9 of DC differentiation were stimulated with LPS (1 μg/ml, 24 hours) and co-cultured with CFSE labeled T cells for 5 days and CD4^+^ and CD8^+^ T cell proliferation was determined by flow cytometry. Representative flow cytometry analysis and quantification of percentage and number of dividing CD4^+^, CD8^+^ and/or CD103^+^ T cells are shown. ConA-stimulated and unstimulated T cells, positive and negative control, respectively. n = 2-3, independent experiments and independent healthy donors; line represents mean; *p < 0.05, **p < 0.005, ***p < 0.001, one-way ANOVA with tukey’s multiple comparisons test versus unstimulated T cells or as indicated.

Taken together, the JAK2 V617F^hom^ DC and the unmutated and healthy control DC obtained here are fully competent in chemotaxis towards CCL19 and in CD4 and CD8 T cells activation. In addition their ability of polarizing CD8^+^ T cells towards CD8^+^ CD103^+^ T_RM_ cells qualifies them as DC3.

## Discussion and Conclusion

DC3 represent a novel CD1c^+^ DC subset recently identified by single cell analysis (Villani et al., 2017; Dutertre et al., 2019). DC3 are unique in their capacity to polarize CD8^+^ T cells into CD8^+^ CD103^+^ T_RM_ cells (Bourdely et al., 2020). Here we describe a DC differentiation system for human DC3 from iPS cells. These iPS cell-derived DC3 have the gene expression and surface marker makeup of blood DC3 and polarize CD8^+^ T cells into CD8^+^ CD103^+^ T_RM_ cells in vitro.

DC3 are within the DC landscape positioned between monocytes and DC2, but resemble more DC2 than monocytes. Additionally, DC3 are proposed to directly develop from DC/macrophage progenitors (MDP) along an IRF8^low^ trajectory and independently of the IRF8high common DC progenitor (CDP) trajectory, which gives rise to cDC1, DC2 and pDC (Cytlak et al., 2020).

iPS cell differentiation in EB protocols into HE and HPC, and then further into mature hematopoietic cells is known to resemble yolk-sac hematopoiesis (Ivanovs et al., 2017; Lee et al., 2018), where only unspecific DC precursors are found and no further DC subsets are distinguished (Popescu et al., 2019). In these protocols myeloid cells, such as primitive macrophages (Takata et al., 2017; Lee et al., 2018), mast cells (Kovarova et al., 2010) and primitive megakaryocytes (Liu et al., 2015), develop rather than definitive lymphocytes (Sturgeon et al., 2014). Thus, in our iPS cell differentiation this myeloid bias in concert with the cytokines Flt3L and IL-7 might favor DC3 differentiation rather than development of other lineages.

Upon DC3 differentiation several DC lineage determining transcription factors, such as *IRF4*, *IRF8* and *PU.1*, were upregulated. Yet the differentiation conditions applied apparently did not support executing bona fide cDC1, DC2 and pDC development. Additionally, whether the DC3 obtained exhibit a primitive imprint as remnant of yolk-sac hematopoiesis is an open question. Thus, an efficient execution of cDC1, DC2 and pDC development might require more definitive hematopoietic progenitors and/or higher IRF8 expression (Sontag et al., 2017; Cytlak et al., 2020).

Whether DC3 development dependents on Flt3L or GM-CSF is controversial (Dutertre et al., 2019; Bourdely et al., 2020). Initial data indicate that in our iPS cell differentiation system GM-CSF caused development of granulocytes and macrophages rather than DC3 (data not shown). In addition, JAK2 V617F^hom^ iPS cells showed a bias towards erythropoiesis and megakaryopoiesis in line with the PV patient phenotype. These data indicate that the differentiation system employed here allows development into further hematopoietic lineages. Interestingly, the JAK2 V617F mutation increased DC3 numbers but did not impact on the DC3 functions analysed so far.

Finally, patients with systemic lupus erythematosus (SLE) show accumulated DC3 in blood, which correlates with SLE disease activity index (Dutertre et al., 2019). In breast cancer patients DC3 infiltration and frequencies of CD8^+^ CD103^+^ T_RM_ cells correlate, which is related to a protective prognosis (Bourdely et al., 2020). Thus, the novel iPS cell differentiation system for DC3 developed here stands as a valuable tool for studying DC3 in human disease and for developing novel therapeutic strategies, such as pharmacologically targeting DC3 in disease.

## Data availability statement

Raw data supporting the conclusions of this article will be made available by authors, without undue reservation.

## Ethics statement

Peripheral blood mononuclear cells of healthy donor and of two PV patients with JAK2 V617F mutation were obtained after informed consent (local ethics board reference numbers EK099/14 and EK206/09).

## Author contributions

T.S.: Conception and design, experiments, collection and assembly of data, data analysis and interpretation, figure preparation and manuscript writing; M.A.S.T., J.B., K.O.: Experiments, collection and/or assembly of iPS cell data, data analysis and interpretation; N.F., K.G.: Experiments, collection and/or assembly of DC data, data analysis, interpretation and figure preparation; C.K., S.S.: Establishment of patient iPS cells; K.S., S.K., T.H.B., N.C.: Data analysis and interpretation; Y.-I.T.: Conception and design; M.Z.: Conception and design, data analysis and interpretation, manuscript writing. All authors approved the final version of the manuscript for submission.

## Acknowledgments

We acknowledge the support of the Interdisciplinary Center for Clinical Research Aachen (IZKF Aachen) FACS Core Facility for cell sorting and the expert administrative assistance of E. Mierau.

## Supplementary Material

**Supplementary Figure 1.**
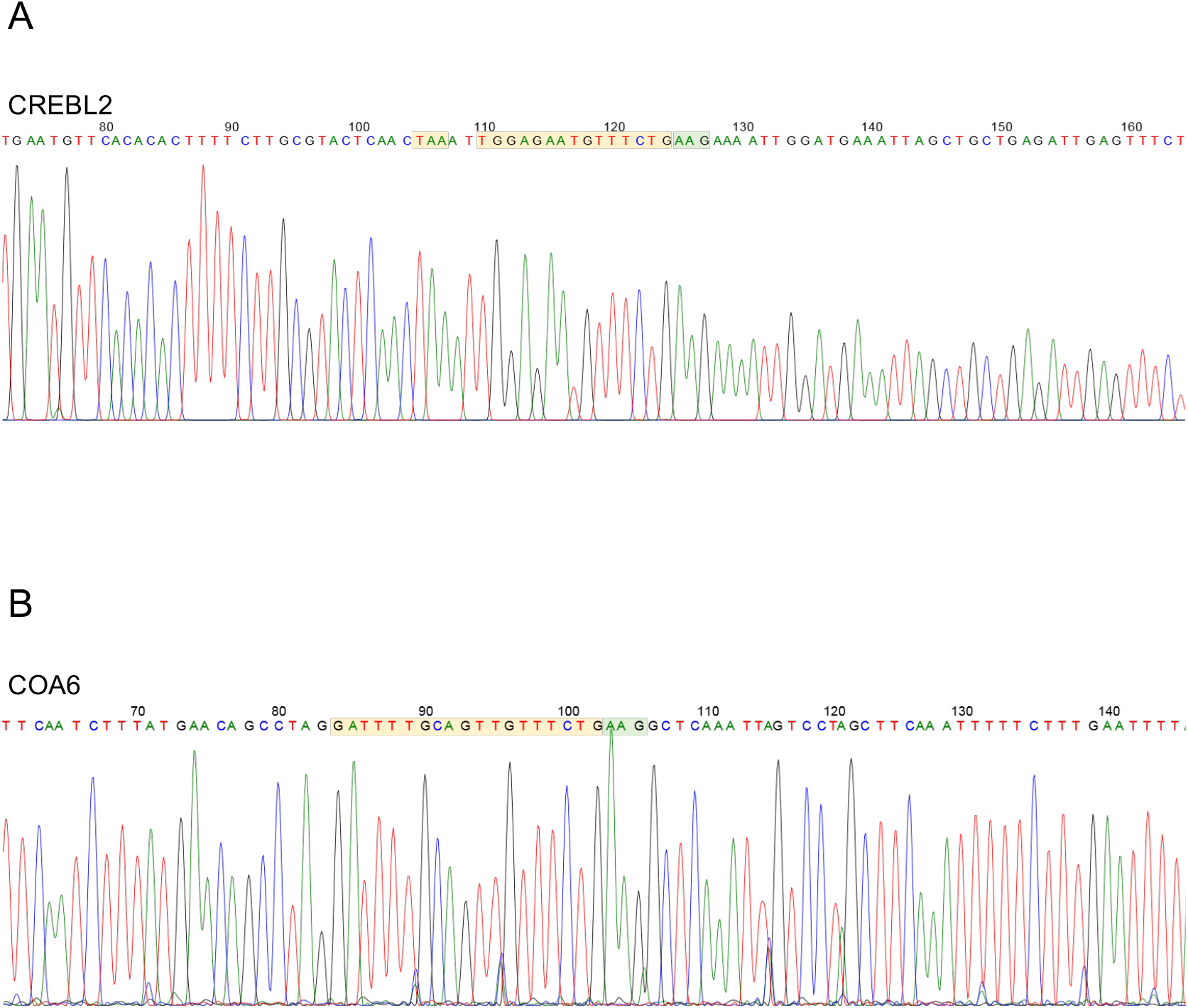
DNA sequence of potential CRISPR/Cas9 off-target genes CREBL2 and COA6. The potential CRISPR/Cas9 off-target genes cAMP responsive element binding protein like 2 (CREBL2) and cytochrome C oxidase assembly factor 6 (COA6) were analysed by DNA sequencing (A and B, respectively) and no off-target effects were found. CRISPR/Cas9 gRNA sequence targeting JAK2 (light orange box); PAM sequence (light green box).

**Supplementary Figure 2.**
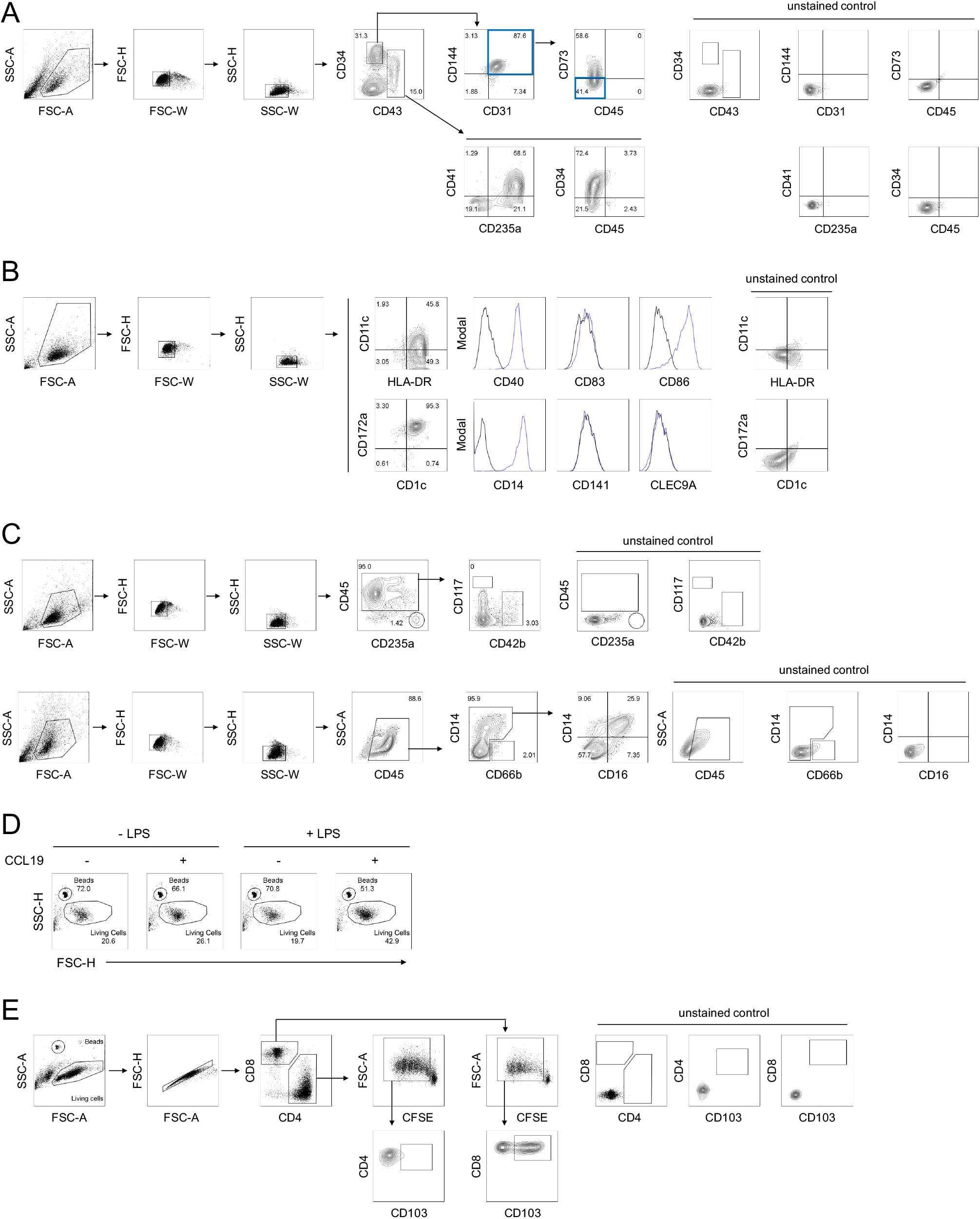
Gating strategies for flow cytometry analysis. **(A)** Gating strategy for CD34+ CD31+ CD144+ CD43- CD45- CD73- HE (blue boxes) and CD43+ CD34^low/-^ HPC in Figure 1B and 2A. **(B)** Gating strategy for HLA-DR^+^ DC and the DC subsets cDC1 (CD141 and CLEC9A) and cDC2 (CD1c and CD172a) and for the co-stimulatory molecules CD40, CD83 and CD86 in Figure 1C, F and G, Figure 2B and C, and Supplementary Figure 1B-D. **(C)** Gating strategy for CD235a/glycophorin A^+^ erythrocytes, CD42b^+^ megakaryocytes and CD14^+^, CD16^+^ and CD66b^+^ myeloid cells (monocytes, makrophages and granulocytes, respectively) in Figure 3A and B. **(D)** Gating strategy for DC migration towards CCL19 chemokine in Figure 4A and Supplementary Figure 2. **(E)** Gating strategy for CD4^+^ and CD8^+^ T cells and for CD103 in T cell activation assays in Figure 4B.

**Supplementary Figure 3.**
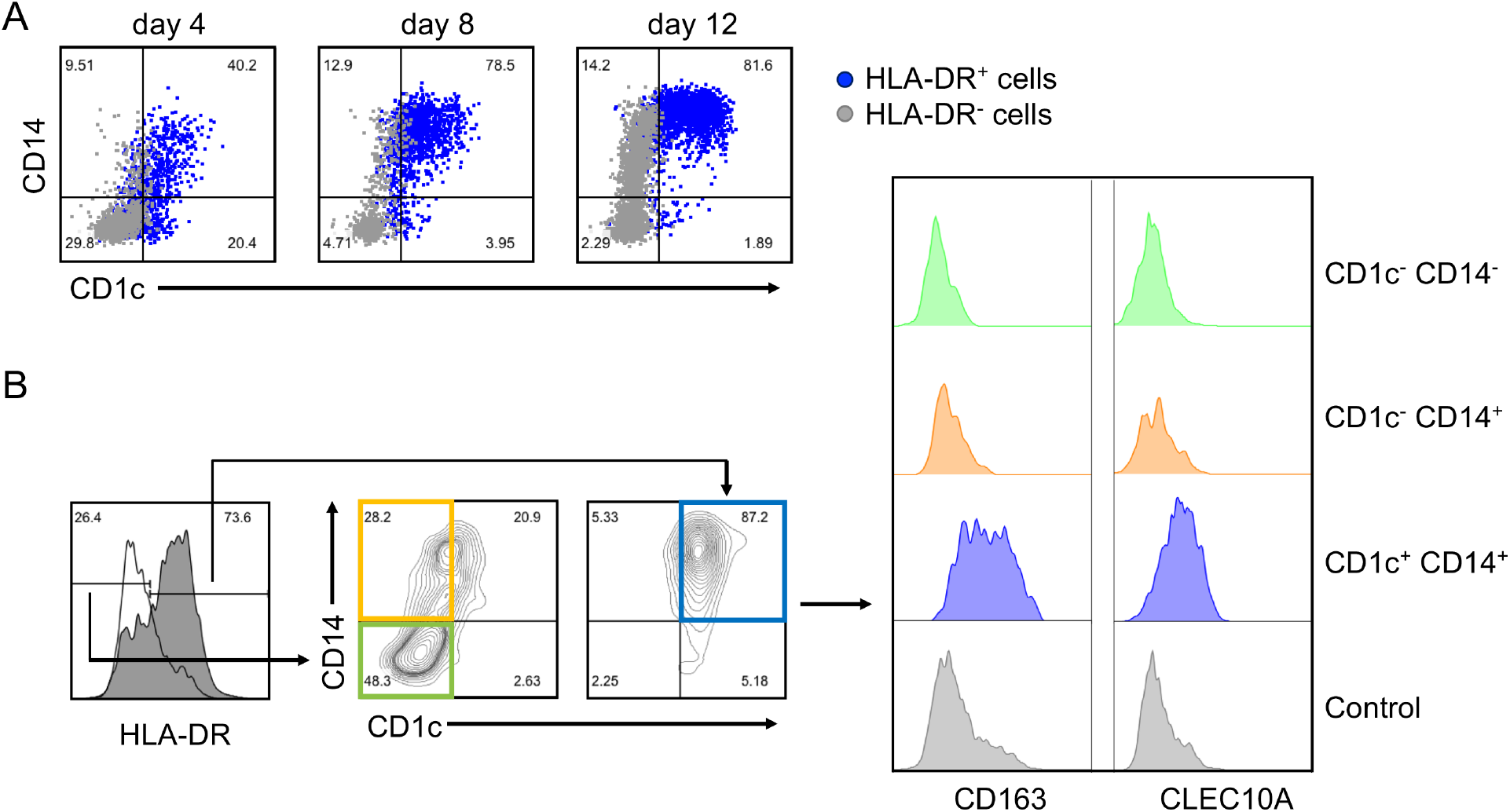
**(A)** Kinetics of emerging HLA-DR^+^ CD1c^+^ CD14^+^ cells of healthy donor during DC differentiation at day 4, 8 and 12 by flow cytometry. **(B)** Representative flow cytometry analysis of CD163 and CLEC10A on HLA-DR^+^ CD1c^+^ CD14^+^ cells of healthy donor on day 7-8 of DC differentiation. Control, unstained cells. *n* = 2

**Supplementary Figure 4.**
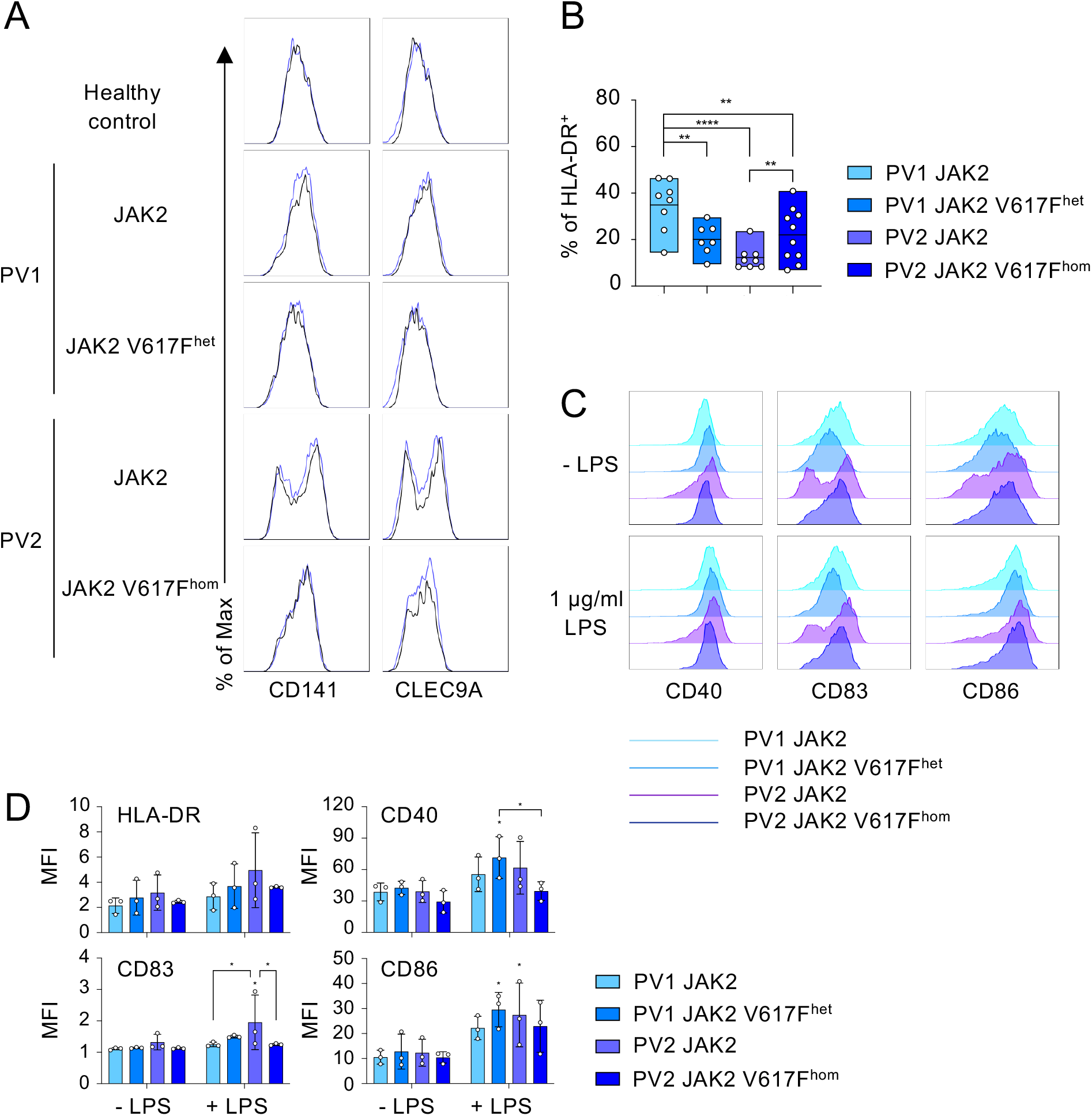
**(A)** Representative flow cytometry analysis of CD141 and CLEC9A on HLA-DR^+^ cells on day 8-9 of DC differentiation. JAK2 genotypes of patients PV1 and PV2, and healthy donor (healthy control) are as in Figure 3A. Blue lines and black lines in histograms represent stained and unstained cells, respectively. *n* = 2-3. **(B)** The percentage of HLA-DR^+^ cells on day 8-9 of DC differentiation for JAK2 V617F^het^ and JAK2 V617F^hom^ cells of patients PV1 and PV2, and for cells without mutation (JAK2). *n* = 7-10, line represents mean; *p < 0.05, **p < 0.005, ****p < 0.0001, one-way ANOVA with uncorrected fisher’s LSD test. **(C)** Representative flow cytometry analysis of CD40, CD83 and CD86 on unstimulated and LPS-stimulated CD1c^+^ HLA-DR^+^ cells on day 8-9 +1 of DC differentiation. JAK2 genotypes of patients PV1 and PV2 are as in (B). *n* = 3. **(D)** Expression of DC activation markers CD40, CD83 and CD86 on unstimulated and LPS-stimulated CD1c^+^ HLA-DR^+^ cells of (C). MFI values were normalized to unstained cells. JAK2 genotypes of patients PV1 and PV2 are as in (B). *n =* 3, mean ± SD; *p < 0.05, two-way ANOVA with uncorrected fisher’s LSD test between unstimulated and LPS-stimulated cells for each JAK2 V617F genotype or as indicated.

**Supplementary Figure 5.**
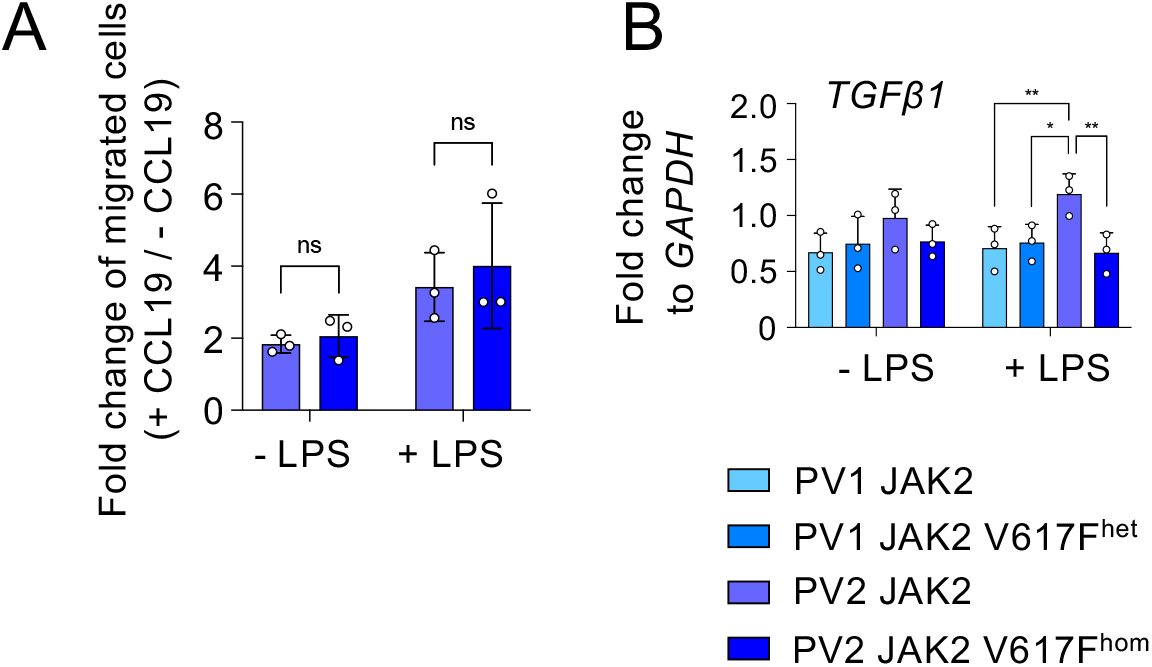
**(A)** Chemotaxis assay towards CCL19 of JAK2 V617F^hom^ and JAK2 control CD1c^+^ HLA-DR^+^ cells as in Figure 4A. Numbers of migrated cells were determined by flow cytometry and the fold change to the cell number without CCL19 is shown. *n* = 3, mean ± SD; ns, not significant. **(B)** TGFβ1 expression in unstimulated and LPS-stimulated HLA-DR^+^ cells of Supplementary Figure 1C determined by RT-qPCR analysis. Values were normalized to *GAPDH* and 2^-ΔCt^ values were shown. *n* = 3, mean ± SD; *p < 0.05, **p < 0.005, one-way ANOVA with uncorrected fisher’s LSD test between unstimulated and LPS-stimulated cells for the various JAK2 V617F genotypes as indicated.

**Supplementary Table 1:**
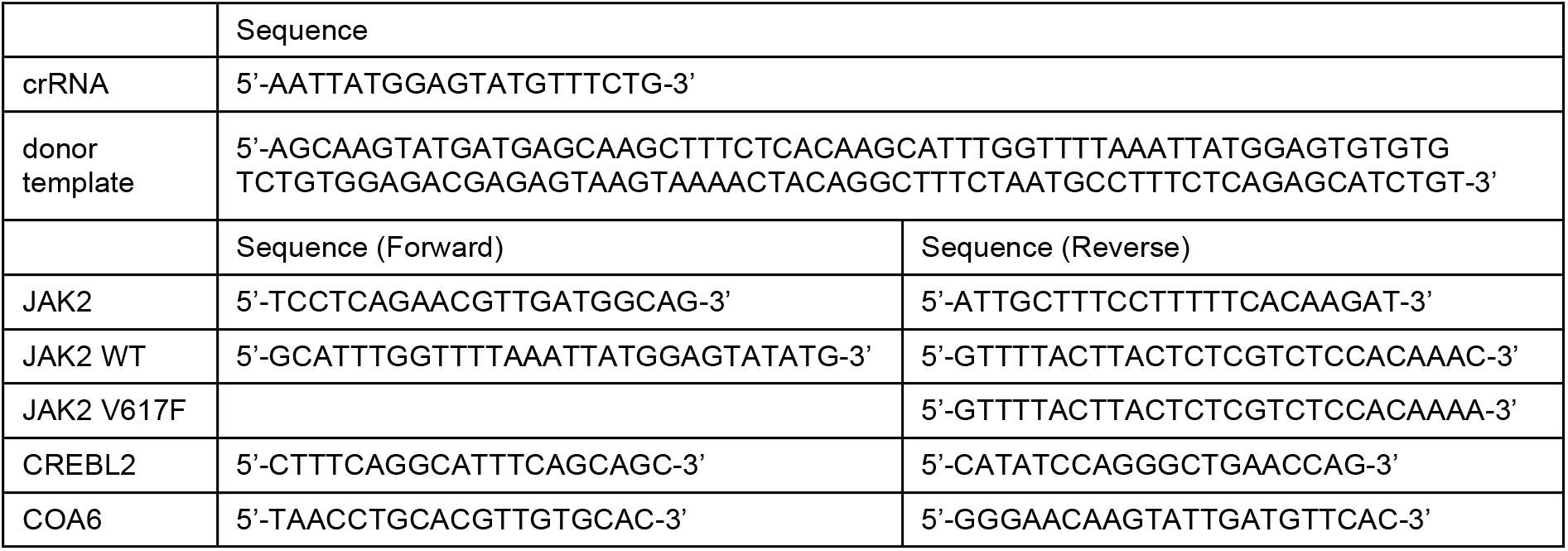
Sequences of crRNA and donor template for CRISPR/Cas9 editing, JAK2 sequencing primers (JAK2) and JAK2 V617F allele specific PCR primers (JAK2 WT and JAK2 V617F) and sequencing primer for CREBL2 and COA6 off-target analysis.

**Supplementary Table 2:**
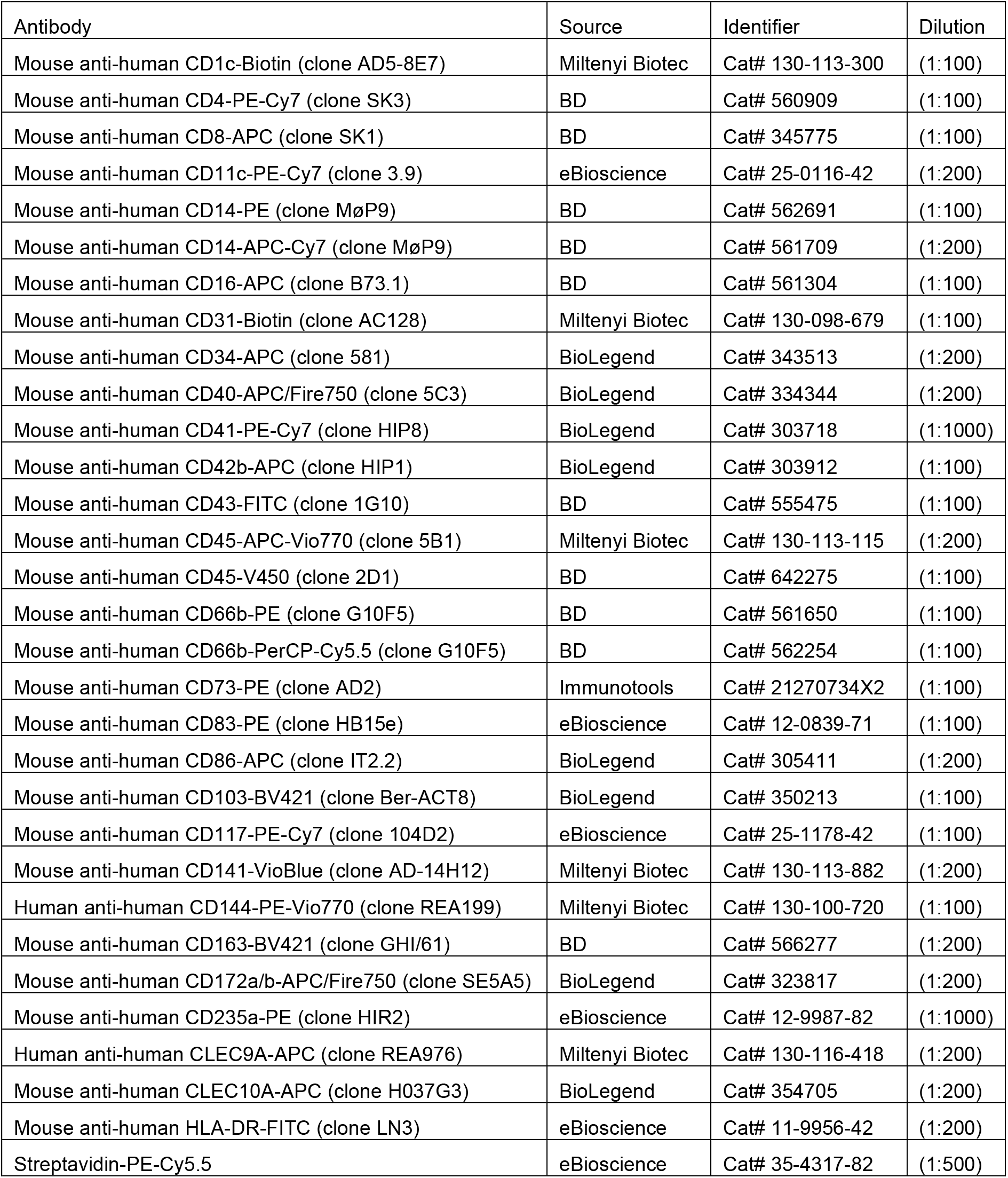
Antibodies used for FACS in this study

| Antibody | Source | Identifier | Dilution |
| --- | --- | --- | --- |
| Mouse anti-human CD1c-Biotin (clone AD5-8E7) | Miltenyi Biotec | Cat# 130-113-300 | (1:100) |
| Mouse anti-human CD4-PE-Cy7 (clone SK3) | BD | Cat# 560909 | (1:100) |
| Mouse anti-human CD8-APC (clone SK1) | BD | Cat# 345775 | (1:100) |
| Mouse anti-human CD11c-PE-Cy7 (clone 3.9) | eBioscience | Cat# 25-0116-42 | (1:200) |
| Mouse anti-human CD14-PE (clone MøP9) | BD | Cat# 562691 | (1:100) |
| Mouse anti-human CD14-APC-Cy7 (clone MøP9) | BD | Cat# 561709 | (1:200) |
| Mouse anti-human CD16-APC (clone B73.1) | BD | Cat# 561304 | (1:100) |
| Mouse anti-human CD31-Biotin (clone AC128) | Miltenyi Biotec | Cat# 130-098-679 | (1:100) |
| Mouse anti-human CD34-APC (clone 581) | BioLegend | Cat# 343513 | (1:200) |
| Mouse anti-human CD40-APC/Fire750 (clone 5C3) | BioLegend | Cat# 334344 | (1:200) |
| Mouse anti-human CD41-PE-Cy7 (clone HIP8) | BioLegend | Cat# 303718 | (1:1000) |
| Mouse anti-human CD42b-APC (clone HIP1) | BioLegend | Cat# 303912 | (1:100) |
| Mouse anti-human CD43-FITC (clone 1G10) | BD | Cat# 555475 | (1:100) |
| Mouse anti-human CD45-APC-Vio770 (clone 5B1) | Miltenyi Biotec | Cat# 130-113-115 | (1:200) |
| Mouse anti-human CD45-V450 (clone 2D1) | BD | Cat# 642275 | (1:100) |
| Mouse anti-human CD66b-PE (clone G10F5) | BD | Cat# 561650 | (1:100) |
| Mouse anti-human CD66b-PerCP-Cy5.5 (clone G10F5) | BD | Cat# 562254 | (1:100) |
| Mouse anti-human CD73-PE (clone AD2) | Immunotools | Cat# 21270734X2 | (1:100) |
| Mouse anti-human CD83-PE (clone HB15e) | eBioscience | Cat# 12-0839-71 | (1:100) |
| Mouse anti-human CD86-APC (clone IT2.2) | BioLegend | Cat# 305411 | (1:200) |
| Mouse anti-human CD103-BV421 (clone Ber-ACT8) | BioLegend | Cat# 350213 | (1:100) |
| Mouse anti-human CD117-PE-Cy7 (clone 104D2) | eBioscience | Cat# 25-1178-42 | (1:100) |
| Mouse anti-human CD141-VioBlue (clone AD-14H12) | Miltenyi Biotec | Cat# 130-113-882 | (1:200) |
| Human anti-human CD144-PE-Vio770 (clone REA199) | Miltenyi Biotec | Cat# 130-100-720 | (1:100) |
| Mouse anti-human CD163-BV421 (clone GHI/61) | BD | Cat# 566277 | (1:200) |
| Mouse anti-human CD172a/b-APC/Fire750 (clone SE5A5) | BioLegend | Cat# 323817 | (1:200) |
| Mouse anti-human CD235a-PE (clone HIR2) | eBioscience | Cat# 12-9987-82 | (1:1000) |
| Human anti-human CLEC9A-APC (clone REA976) | Miltenyi Biotec | Cat# 130-116-418 | (1:200) |
| Mouse anti-human CLEC10A-APC (clone H037G3) | BioLegend | Cat# 354705 | (1:200) |
| Mouse anti-human HLA-DR-FITC (clone LN3) | eBioscience | Cat# 11-9956-42 | (1:200) |
| Streptavidin-PE-Cy5.5 | eBioscience | Cat# 35-4317-82 | (1:500) |

**Supplementary Table 3:**
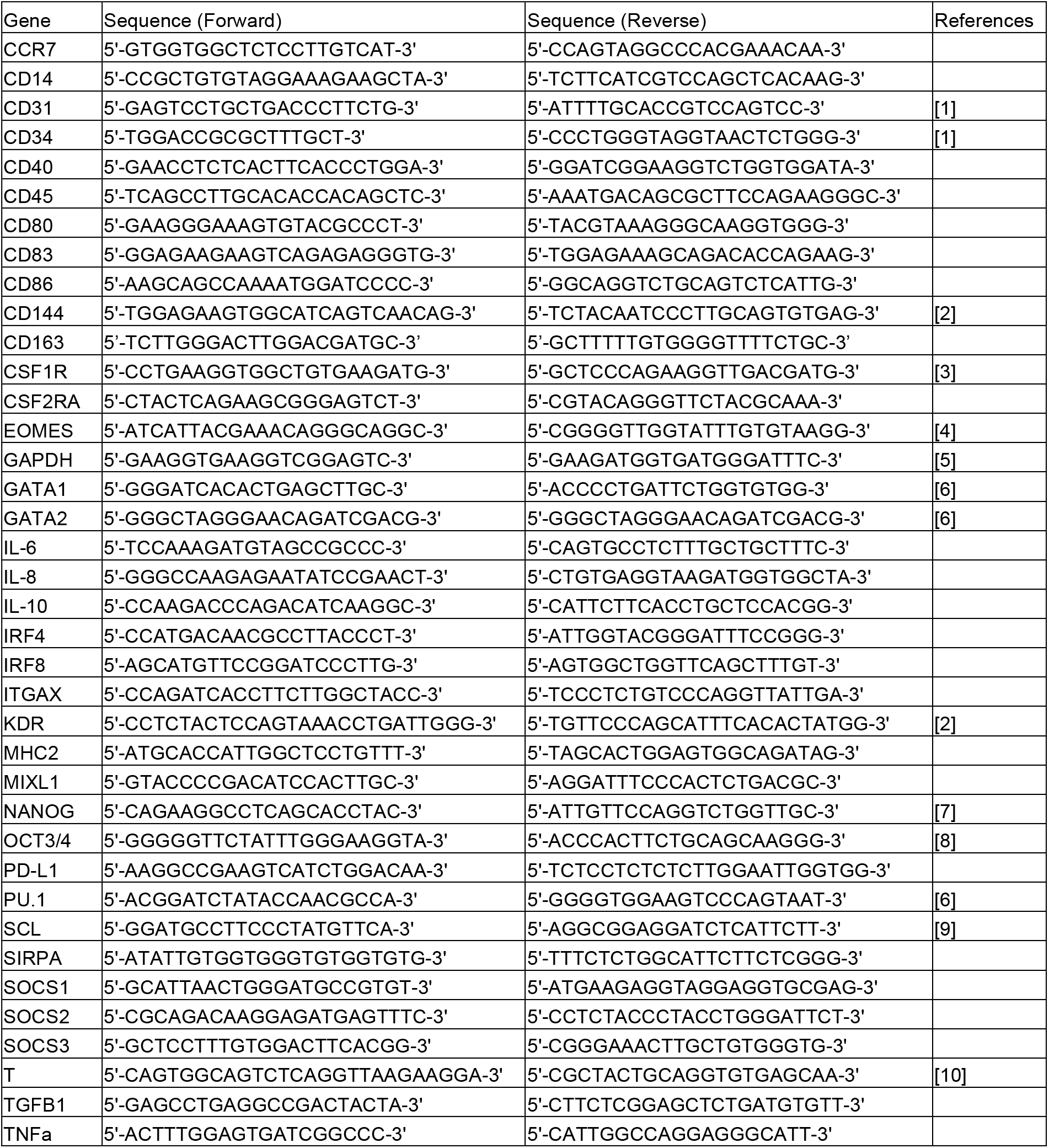
Primers used for RT-qPCR in this study.

